# Spatial and temporal detection of root exudates with a paper-based microfluidic device

**DOI:** 10.1101/2023.12.06.570394

**Authors:** Daniel Patko, Udara Bimendra Gunatilake, Lionel X. Dupuy, Lourdes Basabe-Desmonts, Fernando Benito-Lopez

## Abstract

Root exudates control critical processes in the rhizosphere, retaining water, selecting for beneficial microorganisms or solubilising nutrients prior to uptake by the plant. Analysing root exudation patterns however is challenging because existing methods are often destructive and unable to resolve spatial and temporal variations in the production of root exudates. Here, we present a paper-based microfluidic device with integrated colorimetric sensors for the continuous extraction of root exudates along plant roots. The microfluidic device used standard filter paper wax printer to create channels for water to carry the exudates towards the sensors. TiO_2_ nanotubes/alginate hydrogel-based sensors were used to analyse the glucose content of the root exudates of living plants. The study shows that the paper microfluidic substrate successfully extracts the released glucose from the root, and transfers it to the hydrogel-based sensor to be calorimetrically detected from independent sections of the root at different times, up to 7 days. The method was tested on two different wheat varieties *Triticum aestivum* (rgt Tocayo and Filon varieties), where significant differences in exudation patterns were recorded. The researchdemonstrates the feasibility of low cost technological solution for high precision screening and diagnostic of the biochemical composition of root exudates.

## Introduction

Roots exude biomolecules that modify the physical and chemical properties of their environment. For example, root exudates increase the ability of soils to retain water (Carminati et al., 2010) but also reduce surface tension (Naveed et al., 2019), which is an important characteristic for the rewetting of the soil (Demie et al., 2010; Chaichi et al., 2015). Root exudates contain organic acids that reduce the pH of the soil (Dakora and Phillips, 2002; Jones et al., 2021; Lin et al., 2022), contain enzymes that break down organic materials, contribute to the mineralisation of nutrients (Richardson et al., 2009), and facilitate the solubilisation and transport of mineral elements to the plants (Dakora and Phillips, 2002). Root exudates also contain diverse polysaccharides and compounds with antimicrobial properties such as phenols, peptides, enzymes, amino acids, nucleotides, hormones, organic acids, fatty acids (Baetz and Martinoia, 2014; Steinauer et al., 2016; Sasse et al., 2018; Galloway et al., 2020). These compounds in turn significantly modify microbial communities in the rhizosphere (Narula et al., 2009; Sasse et al., 2018; Ulbrich et al., 2022).

Exudates can be extracted from plants grown in the field, but this requires excavation and washing of the root system. Protocols usually consist of applying gentle wash and several purification steps before the samples can be analysed using conventional analytical techniques such as HPLC and/or mass spectrometry (Oburger and Jones, 2018; Galloway et al., 2020; Williams et al., 2021). To avoid excavation of the root, microdialysis probes have been used to continuously measure the flow of exudates (Plett et al., 2021), but measurement made using such systems are localised and can be difficult to apply into the soil. To overcome the difficulty of working with soil, it is possible to sample liquid solution from hydroponic or semi hydroponic cultures and in turn purify compounds from the extracted solution. Although, exudate extraction in hydroponic systems is easier, they may not correlate well to the exudates produced in natural soil conditions.

Recently, several microfluidic techniques have emerged as powerful approaches to observe and measure rhizosphere processes (Kaiser et al., 2023). For instance, microfluidics have enabled live observation of root interactions with microbes (Massalha et al., 2017; Noirot-Gros et al., 2020; Singh et al., 2021). Artificial soils have been used to control the physical structure of the substrate while enabling live observation within the device (Sharma et al., 2020; Liu et al., 2021). A growing number of studies have successfully demonstrated the potential of microfluidic technology for the fine-tuning and analysis of plant growth (Grossmann et al., 2011; Parashar and Pandey, 2011; Stanley et al., 2016; Stanley and van der Heijden, 2017; Aufrecht et al., 2022). Nevertheless, there are several factors preventing their wide application in plant science, such as the need of special equipment, the elevated costs, and the time required for the prototyping and the fabrication of the devices.

In many cases, photolithography techniques (Mitra and Chakraborty, 2012; Aufrecht et al., 2022; Kaiser et al., 2023) use light sensitive material such as SU-8 or Norland optical adhesive to create the moulds to imprint various materials (Mitra and Chakraborty, 2012; Scott and Ali, 2021). 3D printing can significantly decrease the fabrication time of microfluidic devices, but the costs remain high and the choice of materials is limited (biocompatibility) (Sanchez Noriega et al., 2021; Kaiser et al., 2023). In other cases, the combination of porous membranes with conventional microfluidics allows sampling root exudates at different places of the root trough time (Patabadige et al., 2019). Microfabrication techniques can be employed to mimic the structure of soil (Aufrecht et al., 2022; Walton et al., 2022) or the mapping of simple chemical changes like pH within the rhizosphere (Rudolph-Mohr et al., 2017; Jones et al., 2021; Patko et al., 2023), but the detection of specific molecules remain challenging.

Attempts at developing paper microfluidic devices for application in plant sciences have emerged recently. Applications have focused on developing soil sensors to detect pollution, nutrient composition or moisture content (Aeby et al., 2023). The development of plant wearables, where the microfluidic device is directly applied to the living plant, have also been applied to extract and quantify various analytes (Yin et al., 2021). Recently a filter paper-based colorimetric sensor was presented to detect volatile compounds of stressed plant samples within a Petri-dish; however, the paper was not used as a fluidic element, just as a sensor substrate (Zedler et al., 2023), including electrochemical sensors. The passive capillary forces within the paper has been used to deliver soil pollutants like heavy metals to the electrochemical detection device (Mentele et al., 2012; Rattanarat et al., 2014).

In most previous fluidic devices, paper was viewed and used as substrate for the sensor but not for the plant itself. In this study, we propose the development of a paper-based microfluidic device to extract root exudates from precise locations along the root at regular time intervals. In order to do that, wax patterns were printed using commercial a printer (Carrilho et al., 2009; Lind et al., 2021) to create well defined fluidic channels able to direct the flow of liquid with the exudates though the paper (Potter et al., 2019; Catalan-Carrio et al., 2021). We used the technique to print vertical channels that transport exudates towards a TiO_2_ nanotubes (TNT)/alginate hydrogel-based glucose sensor polymerised at the outlet of the paper channel (Gunatilake et al., 2021). The exudate profiles of two wheat varieties (*Triticum aestivum* var. rgt Tocayo and Filon) were analysed, demonstrating the ability of this microfluidics technology to detect phenotypic differences in root exudation profiles over time and at different positions of the root.

## Materials and Methods

### Fabrication of the paper-based microfluidic device

The developed device substrate was based on Whatman Grade 1 filter paper, 180 μm thickness, (1001-917, Sigma Aldrich, Spain). The designs were drawn in Corel Draw Home and Student Suit X7 (Cleverbridge, Spain) and were printed out using a Xerox ColorQube 8580 printer. A laser cutter (VLS2.30DT, Universal Laser Systems, Spain) was used to cut out the paper to precise dimensions, then the wax was melted into the paper using a hotplate at 120 °C for 30 s. The patterned paper substrate was placed between a microscope slide (631-1550, WVR, Spain) and a 26 x 76 x 2 mm slide of polydimethylsiloxane (PDMS) (Sylgard 184 silicone elastomer 1:10 kit, Farnell, Spain). For the assembly of the device, the surfaces of the glass slide and the PDMS slide were activated by oxygen plasma (PDC-002-CE, Harrick plasma, USA) for 30 s (Mitra and Chakraborty, 2012). After activation, the paper channels were placed onto the glass slide before placing the PDMS slide on top of it. The glass and the PDMS covalently bonded to each other between the inlets and outlets after gentle pressure, creating a microfluidic system with several parallel horizontal channels (Ch) (Figure 1A & B).

**Figure 1.**
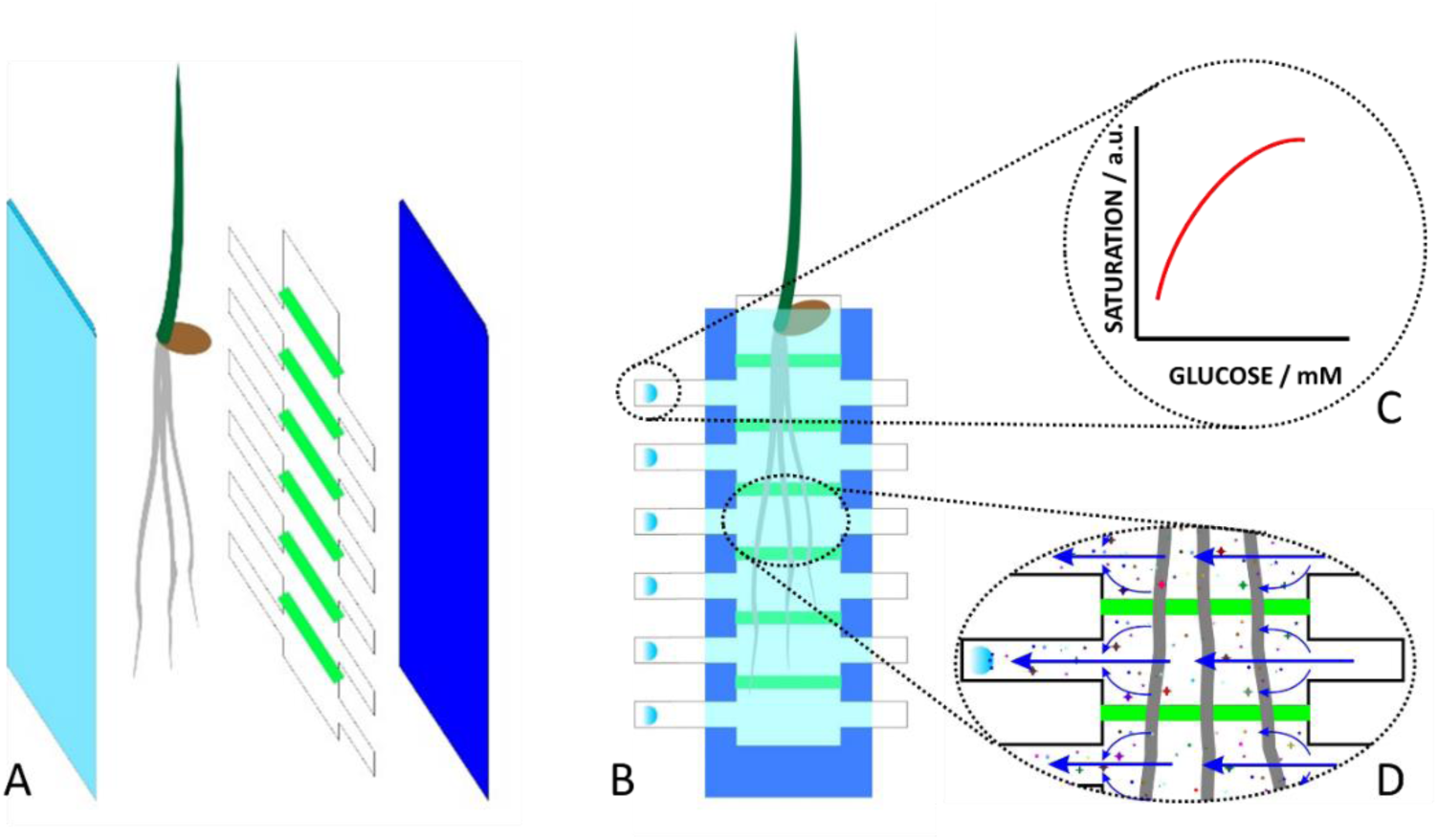
Schematic representation describing the assembly of the wax printed paper-based microfluidic device and its working principle. A) Filter paper is laser cut to produce de shape of the microfluidic device with a 6 inlets (right of the chip) and 6 outlets (left of the chip). Wax patterns (green) are printed on the paper to direct the flow towards the sensor located at the outlet. The paper-based microfluidic device is placed into a chamber (consists of a glass and a PDMS slide) so that plant roots can grow healthily for few days in contact with the paper substrate. B) A TNT/alginate hydrogel-based glucose sensor is placed on the paper at the outlet (blue spot on the left side of the device). C) Image analysis allows automatic quantification of the state of the sensor. D) Growing roots deposit biomolecules on the paper, which dissolve in water and move with the water flow to the outlet of paper device, where the sensor is located. The wax patterns (green) separate the different areas from the root and transport the biomolecules towards the sensors, from right to left (blue arrows).

The design used in this study consisted of 8.0 mm wide channels separated by 2.0 mm wide printed barriers. The width of the channels was chosen so that a sufficient quantity of exudates from the roots were collected and detected by the sensor. It also facilitates the acquisition of image data from the sensors and allows sampling the paper itself for further analysis, for example using conventional analytical techniques. The inlets (right hand side of the paper) were 4.0 mm wide and 10.4 mm long, and the outlets (left hand side of the paper) were 4.0 mm wide and 15.8 mm long. The longer length of the outlets improved the speed of glucose extraction because evaporation increases with the surface area of the paper substrate. Details of the assembly and the designs are presented in the Supplementary Information SI-1.

### Optimisation and testing of the paper channels

In order to understand the flow behaviour in the channels, yellow and blue food dyes (Sabadell, Vahiné, Spain) were used to visualise the flow. The dyes were diluted 10 times in distilled water before flow study. 5.0 µL of the solution were pipetted in the inlet, pipetting was repeated to maintain saturation of the paper until the liquid reached the outlet part of the channels (Figure 2 A). We tested the permeability of the wax strip of 1.0 mm, 1.5 mm and 2.0 mm width (Supplementary Information SI-2). The flow speed was determined based on the position of the colour front at different time points. All tests were carried out in the presence of the plant roots.

**Figure 2.**
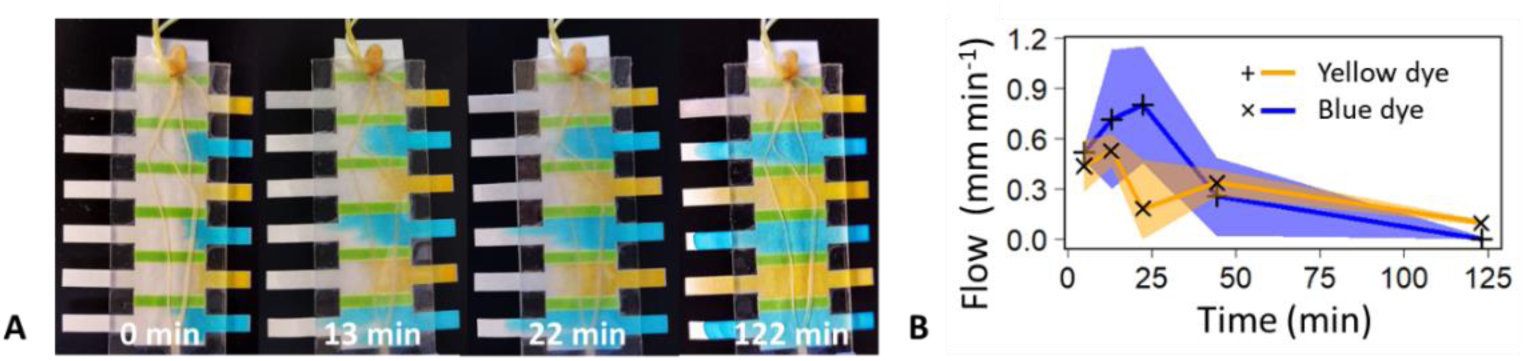
Flow characterisation in the paper-based microfluidic device. A) Pictures of the paper-based microfluidic device at different times during flow experiments using blue and yellow dyes, showing the evolution of the flow, from left to right. B) Graphic representing the flow speed as a function of time of the dyes. Shaded areas correspond to the standard deviation of the flow speed for each dye (n = 3).

### Plants used in this study

Two varieties of winter wheat *Triticum aestivum* (var. rgt Tocayo and var. Filon), were used for the experiments. All seeds were surface sterilised according to O’Callaghan et al. (2018). The seeds were soaked in distilled water for 1 h, and then immersed into 2 % of calcium hypochlorite (211389, Sigma-Aldrich, Spain) solution for 15 min. Finally, the seeds were rinsed thoroughly using 50.0 mL distilled water for 10 times. The sterilised seeds were placed onto 0.5 % water agar gel plates (J10654, Thermo Scientific, Spain) wrapped in aluminium foil at 22 °C for one day. The seedlings were inserted into the paper-based microfluidic devices and were incubated for 6 days allowing the roots to reach the bottom channel of the device.

The samples were kept in a plastic box containing a wet filter paper to maintain constant humidity inside the box. Plants were kept at 22 °C under 14 h light and 10 h dark periods. In order to provide enough replicates, four experimental runs with five replicates were performed with each of the wheat varieties. Each run included two samples without plants as control. Therefore, the total number of samples for each of the wheat varieties was 20 and the total number of the control samples was 8.

### TNT/alginate hydrogel-based glucose sensor fabrication

The TNT/alginate hydrogel-based glucose sensor solution was done following a similar protocol as the one described by Gunatilake and coworkers (Gunatilake *et al*., 2021). Then, the sensor solution was polymerised on the top of the paper at the outlet of the paper channels (left side of the channel). The sensors were prepared just before inserting the seeds into the microfluidic devices to ensure uniform water content across the channels. The initial colour of the sensor is white and the presence of glucose changes its colour to blue, with the colour intensity related to the glucose concentration. The enzyme complex used for detection consisted of 0.40 mg mL^-1^ glucose oxidase (G2133, Sigma-Aldrich, Spain), 0.05 mg mL^-1^ horseradish peroxidase (77332, Sigma-Aldrich, Spain) and 3,3′,5,5′-tetramethylbenzidine (TMB) (860336, Sigma-Aldrich, Spain) dissolved in dimethyl sulfoxide (> 99.7 %, Sigma-Aldrich, Spain) (24.0 mg : 2.25 mL). The enzyme complex was encapsulated in a sodium-alginate (W2011502, Sigma-Aldrich, Spain) hydrogel matrix with additional TNTs (synthesised according to the publication from Gunatilake and collaborators (Gunatilake et al., 2021)). The TNT content provided a bright white colour to the hydrogel, improving the observation of the colour changes due to a better contrast of the material. All without the need of modifying the enzymatic reaction and concentration of reagents. More information on the preparation of the sensor can be found in the Supplementary Information SI-2.

### TNT/alginate hydrogel-based glucose sensor calibration

For the calibration of the sensor little wax circles were printed (1.0 mm line width, ID = 3.0 mm) on paper, and 2.0 µL of the hydrogel solution with the enzymatic assay was pipetted into each of the circles, Supporting Information SI-4, SI-Figure 3A. In the next step, in order to initiate the polymerisation of the alginate, the filter paper was immersed into 0.4 M CaCl_2_ solution for 1 min. Then, the paper with the polymerised sensor was rinsed in distilled water for 1 min and let to dry (SI-Figure 3B). The devices were fabricated just before the control measurements. SI-Figure 3C shows a picture of a set of glucose sensors in the circular wax printed-paper configuration. The blue colour corresponds to sensors that have detected glucose.

**Figure 3.**
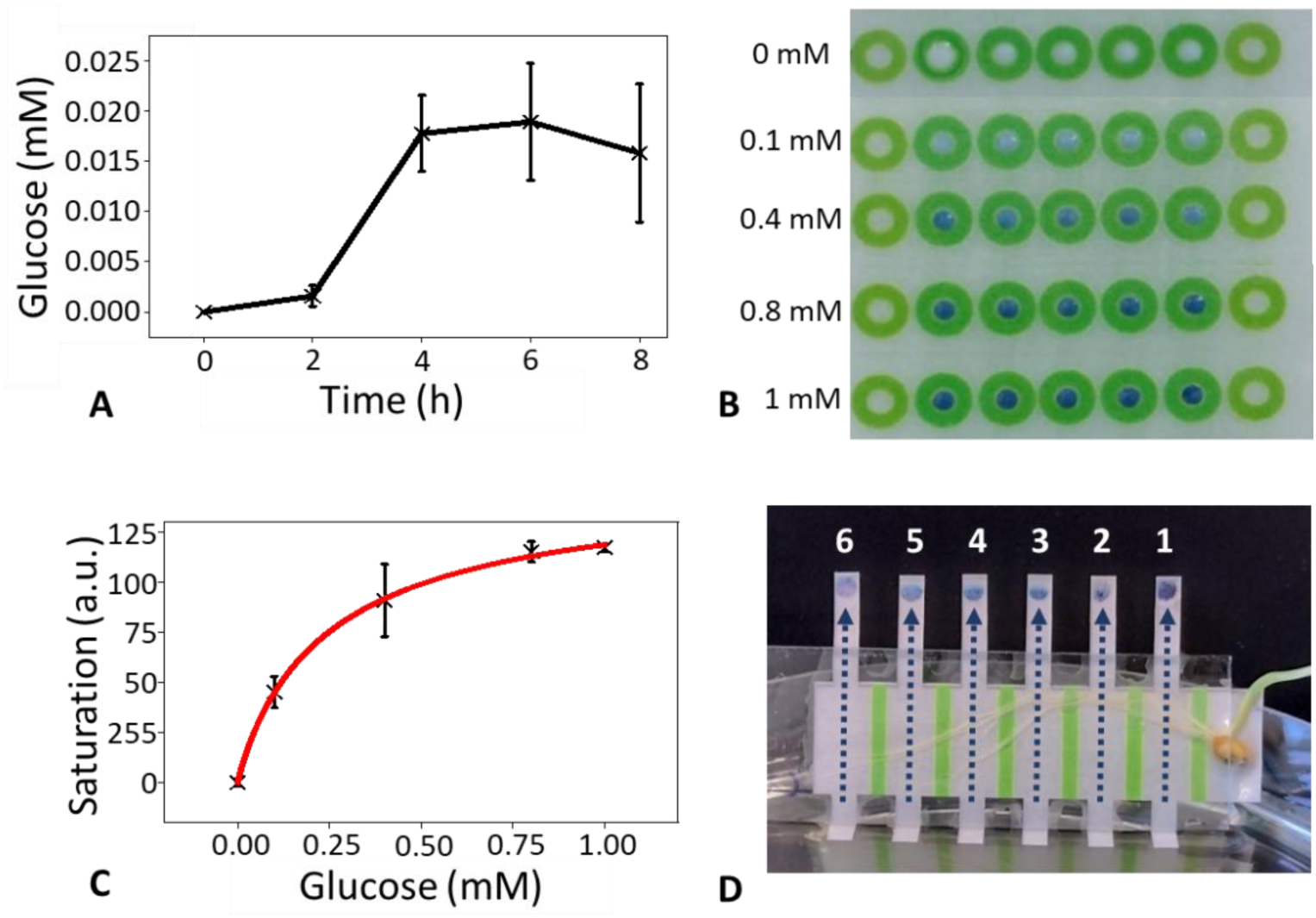
Characterisation of the TNT/alginate hydrogel-based glucose sensor. A) Figure presenting the increase of concentration of glucose over time measured with the sensor in the paper-based microfluidic device under a continuous flow of glucose (1.0 mM). Signals produced by the sensor as a function of time indicated the time needed for all available glucose accumulated in the paper to reach the sensor until saturation. B) Pictures of discrete TNT/alginate hydrogel-based glucose sensors in paper, at different glucose concentrations for calibration. Darker blue colour represents higher glucose concentration. C) Calibration curve using the Hill-equation fitted from the image saturation data and the concentration of glucose. Error bars indicate the standard deviation (n = 5). D) Photograph of a paper-based microfluidic device with inlets immersed in water. The position of the device triggers the flow direction through the channel (dashed arrows) which permits collection of the exudate coming from the plant and subsequent transportation of the exudates to the TNT/alginate hydrogel-based glucose sensor present at the outlet of the device.

A similar protocol was used for the glucose sensor deposition in the microfluidic chip. In this case, the sensors were placed into the left end side of each of the paper-based microfluidic device outlets without any wax encirclement.

Then, glucose solutions were prepared in distilled water at a concentration of 0.0, 0.1, 0.4, 0.8 and 1.0 mM (G8270, Sigma Aldrich, Spain). 2.0 µL of the glucose solution were pipetted on each of the sensors (n = 5). For the control, 2.0 µL of distilled water were used instead of the glucose solutions. To keep the sensors humid, during the chemical reaction, the paper strips were kept in a sealed Petri-dish and signals from the sensors were captured after 90 min.

Images were captured every 15 min and transformed to HSB images (Hue, Saturation and Brightness) using the image processing program ImageJ (version 1.53t 24 August 2022, NIH) (ImageJ). The saturation images were used to identify the region of interest, and the mean saturation values from the circular areas were used as glucose concentration value from the sensor (see Supplementary Information SI-5). The relationship between the mean saturation value and the glucose concentration was fitted by the Hill-equation and used to quantify glucose content from the exudates of the roots. Since the glucose solution covered the whole surface of the sensors, the whole surface of the sensor was used for the calibration.

### Glucose extraction from the microfluidic devices

After 6 days of incubation, the devices containing the plants were placed into Petri-dishes (391-0565, VWR, Spain) filled with 40 mL of distilled water. The inlets (right end side) were immersed into the water, while the outlets faced upward outside of water. Water evaporation at the outlet side of the device created a gradient in water content, which resulted in a flow of water and exudates to the sensor by capillary action from the inlets that where immersed in the water.

The time needed for the glucose accumulated in the paper to move through the channel and to be detected by the sensor was evaluated. In order to do that, 2.0 mm diameter holes were punched into the PDMS (2.0 mm, Uni-Core punch, Harris, Spain) at the middle of the channels. Then, 1.0 µL of 1.0 mM glucose solution was pipetted into each of the channels. Finally, the glucose was extracted using the evaporation process explained above. The glucose signals were captured every 2 h for 8 h.

For the experiments to measure the concentration of glucose in the root exudates, a continuous flow through the device was maintained for 2 h before measurement. Images of the sensors were captured with a digital camera (DS-FI3, Nikon) using a stereoscope (NSZ-810, Balea Optic, Spain) and analysed using ImageJ. Once the images were captured, the devices were placed back to the growth box (see Supplementary Information SI-6). The colorimetric signals from the six sensors were daily collected for seven days and individually measured. The different sensors were numbered from 1 to 6 respectively from top to bottom.

## Results

### Fabrication of the wax printed patterned paper-based microfluidic device

Different prototypes were tested in this study. In general, the design for the analysis of root exudates consisted of a microscope slide, a wax printed filter paper-based microfluidic sheet and a PDMS slide (Figure 1A). In particular, the PDMS slide, which covers the paper channels, was a fundamental part of the device since it is a gas permeable but impermeable to water layer, ensuring a humid and aerated environment for the root to grow. The channels in the paper layer were generated by laser cutting and the wax stripes (green) were printed onto the paper in order to define discrete areas for analysis. As presented in the experimental section, the size and shape of the printed wax stripes affected the flow of liquids through the paper channel. For instance, thin wax stripes (< 2.0 mm) were not impermeable and allowed parallels flows to mix between the different channels. On the other hand, thick wax stripes can (> 2.0 mm) occupy large areas on the paper sheet limiting the number of sensors in the final paper-based microfluidic device configuration (see Supporting Information SI-2). Wax stripes of 2.0 mm width were found to be impermeable for the whole duration of the experiments and were used in the final device.

### Flow characterisation of the paper-based microfluidic device

Food dyes initially flowed at 0.52 ± 0.05 mm min^-1^ in case of the blue dye, and a little bit slower in the case of the yellow dye solutions 0.44 ± 0.02 mm min^-1^ (Figure 2). Nevertheless, 50 min minutes after application of the dye, the speed was reduced and stabilised at 0.25 ± 0.2 mm min^-1^ with the blue dye and 0.33 ± 0.08 mm min^-1^ with the yellow dye. The time for the two food dyes to reach the sensing zone of the channels was approximately 35 ± 10 min.

### Extraction of the accumulated glucose present in the paper channels

We ran a simple experiment to determine the time used to carry out a measurement of the glucose concentration. 1 mM glucose solution was deposited at the inlet (Figure 3A) and the response of the sensor was monitored through time. It took 4 h for the sensor to be saturated. Reducing the time to perform a measurement would therefore reduce the chances for the sensor to be saturated and induce less gravitropic stimulation of the plant. However, reducing time excessively would also increase the limit of detection. Based on this results, we chose to fix the time of extraction of the exudate to 2 h which represents a compromise between keeping the plant growing in vertical position most of the duration of the experiment, and increasing the limit of detection of the sensor.

### TNT/alginate hydrogel-based glucose sensor calibration

The calibration of the glucose sensor was performed using different concentration of glucose solutions, and results show the saturation value increased consistently with glucose concentration (Figure 3B). The relationship between glucose concentration and image saturation was nonlinear and exhibited a plateau for glucose concentration exceeding 1 mM at a saturation value of 117 ± 2 a.u (Figure 3C). The relationship was captured by a Hill-equation (Hill et al., 1910) y = V_max_ x^n^/(k^n^+x^n^), where V_max_ = 151.71 a.u., k = 0.255 mM, n = 0.929, R^2^ = 0.9956, and x is the glucose concentration (mM) and y the saturation value in the image (a.u.). The calibration curve allowed the determination of the glucose concentration exuded by the wheat roots (Figure 3C). In the control (0 mM of glucose), the signal recorded was equivalent to 0.004 mM glucose. The limit of detection (LOD) was calculated as three times the standard deviation (SD) of the values measured in control samples, and the limit of quantification (LOQ) was calculated as ten times the SD of the values detected in control. Using these calculations, we obtained a LOD of 0.012 mM and a LOQ of 0.04 mM, respectively.

### Determination of glucose concentration on different wheat varieties along the roots

Two wheat varieties, *Triticum aestivum* var. rgt Tocayo and var. Filon, were investigated in order to demonstrate the capability of the paper-based microfluidic device to distinguish exudation patterns from different varieties and at different positions along the root. The two varieties grew in the paper-based microfluidic device for 7 days and did not show signs of stress for the entire duration of the experiment. Moreover, there was no observable growth differences between the two wheat varieties with mean root lengths for Filon and rgt Tocayo measured at 51 ± 2 mm and 49 ± 21 mm respectively (p = 0.73).

During the experiments (Figure 3D), changes in the coloration of the sensors were observed over time (Figure 4). At the beginning of the experiment, the coloration of the sensors was observed at the base of the root system, close to the seed, at the right-hand side of the sensor spot (SI-Figure 4A). Over time, the sensors on the top became darker and the sensors at the bottom of the device started to change colour.

**Figure 4.**
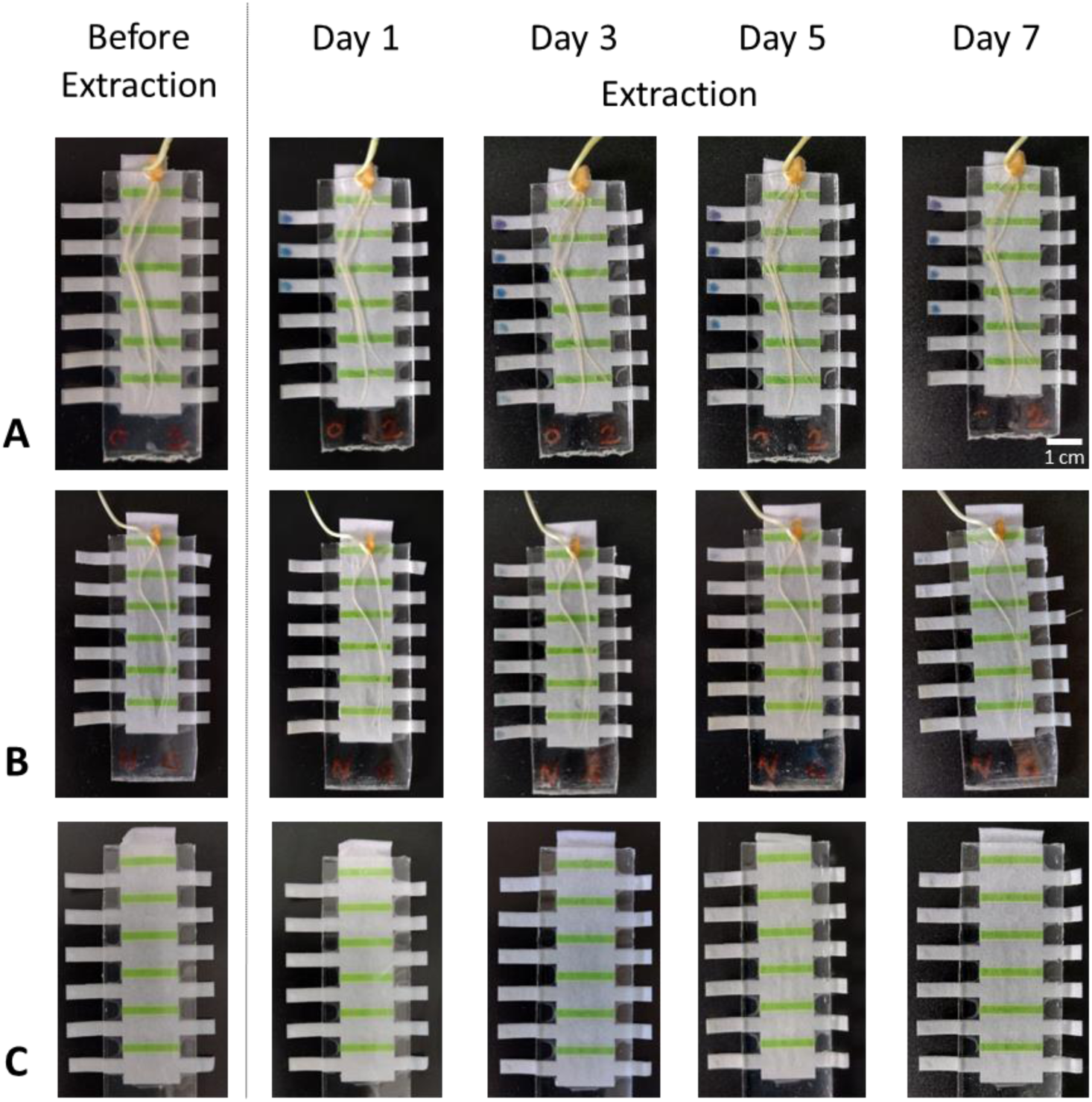
Pictures of the paper-based microfluidic device before and after collection of the glucose exudate from the root for the two wheat varieties as a function of time, up to 7 days. **A)** Paper-based microfluidic device for glucose exudation determination of rgt Tpcayo variety. **B)** Paper-based microfluidic device for glucose exudation determination of Filon variety. **C**) Paper-based microfluidic device without plant, control experiment.

The two wheat varieties had different exudation profiles. Glucose was first detected from the rgt Tocayo variety at sensor positions 1, 2 and 3 during the first days of the experiments (Figure 4A) and then, the detection of glucose was observed for the positions downward (4, 5 and 6). On the other hand, the Filon variety (Figure 4B) did not get substantial coloration of the sensors until day 4 and mainly in the positions 1, 2 and 3 of the paper-based microfluidic device. Finally, the control experiments (Figure 4C) did not present any significant colouration of the sensors.

The quantitative analysis of the signals recorded on the sensors confirmed visual observations (Figure 5 A and B). The distribution of the glucose concentration collected by the sensor at different positions of the root and times indicated that the accumulated amount of glucose is higher in the mature part of the root system within the first few days of the experiment. The lower root parts (lower channels) reach their maximum in exudation later in time (day 3 for the rgt Tocayo and day 5-6 for the Filon).

**Figure 5.**
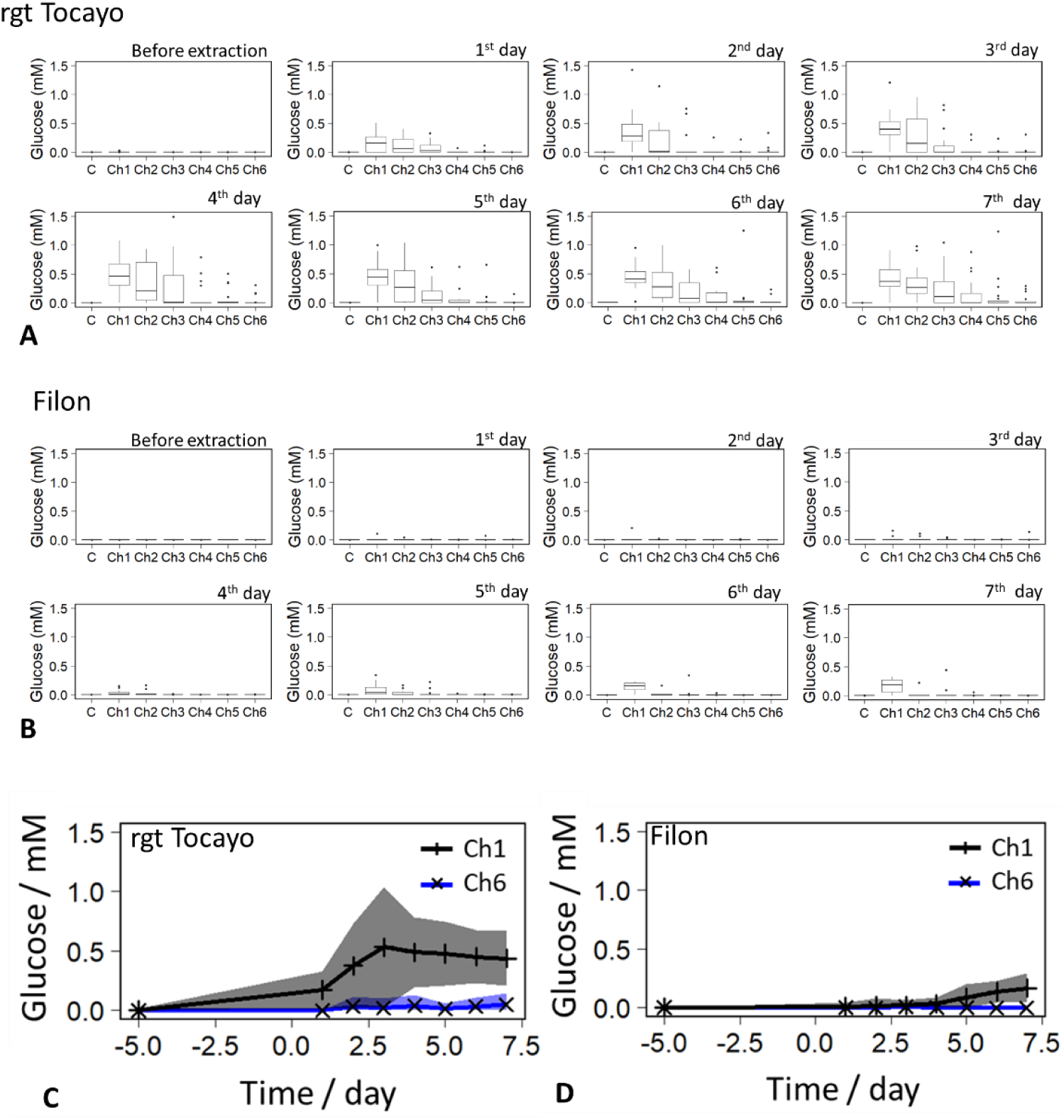
Graphical representation of the glucose concentration in root exudates as a function of time and position along the root. A) Glucose concentration values obtained in the TNT/alginate hydrogel-based glucose sensor for the different channels (Ch), over time, for the rgt Tocayo variety. B) Glucose concentration values obtained in the TNT/alginate hydrogel-based glucose sensor for the different channels, over time, for the Filon variety. The value of the control measurements is indicated with C. The error bars show the standard deviation (n = 20). C) Accumulated exudation of rgt Tocayo variety as a function of time for the first and sixth channels. The grey and the blue shadows correspond to the standard deviation of the saturation; n = 20 and n = 17, respectively. D) Accumulated exudation of Filon variety as a function of time for the first and sixth channels. The grey and the blue shadows correspond to the standard deviation of the saturation; n = 12 and n = 16, respectively.

Moreover, the concentration of exuded glucose was significantly different between the two varieties (see SI-Table 1 and 2 for each time point and each channel). Experiments using rgt Tocayo variety showed that at the base of the root system where the concentration of glucose was the highest, the rate of increase of glucose productions was on average 0.035 ± 0.121 mM, which correspond to about 272 ± 942 ng of glucose, per day resulting in more than 0.5 mM glucose deposited on the sensor during the first 4 days of growth. It was observed that some of the samples recorded a total amount of deposited glucose of 1 mM approaching to the saturation zone of the sensor. In the case of the Filon variety, a very low exudation activity was observed at the beginning of the experiment. The exudation started to increase from day 5 and 6, but it was only significant in the first channel (p = 0.023), with a maximum production of glucose of about 0.003 ± 0.037 mM per day.

## Discussion

### Paper microfluidics for plant science research applications

Unlike other materials used commonly in microfabrication and microfluidic research (PDMS, glass or plastics), paper has a long history of application in plant germination studies, with work on germination paper published as early as a century ago (Morinaga, 1926) Papers have become ubiquitous in root and germination studies because they are hydrophilic and conduct water easily which facilitates watering and use of nutrient solution. Paper is also a good proxy for soil since it retains water and air within its pores, which allows watering and nutrition of the plant without continuous fertigation. Paper is also available at a range of pore size and thickness, and their properties are repeatable and well-studied. Its use in to study root or root-microbe interactions is still very common, for example in the so-called pouch system where the germination paper is placed in between films that limits water loss by evaporation (Collins et al., 2013).

Germination paper is also central to more sophisticated high-throughput plant phenotyping systems. It has a low cost and is suitable to large-scale studies (Atkinson et al., 2015; Dupuy et al., 2017). The root system of plants growing on paper developed in a 2D plane facilitates stacking (Le Marié et al., 2014) and the application of high resolution and low-cost imaging systems such as cameras and flatbed scanners for data acquisition (Adu et al., 2014; Canto et al., 2018). They are generally available in a range of colours, which facilitated automated processing and analysis of image (Pound et al., 2013, 2017; Adu et al., 2016).

Paper is also used in less usual experimental systems, for example as containers or as a wig to deliver nutrient solutions to the roots (Di Salvatore et al., 2008; Adu et al., 2014). Recently, wax printed papers were even used as an experimental technique to affect water availability to growing plants (Lind et al., 2021). Papers are also known to be suitable to transport various molecules reliably since it has been commonly used in sensors (Mentele et al., 2012; Rattanarat et al., 2014). It is clear, therefore, that paper microfluidic technologies could easily be adopted by the plant science community.

### Detection of root exudates using paper microfluidic system

We have developed a hybrid patterned filter paper-based microfluidic device to extract and quantify how root exudation varies spatially and temporally. The configuration of the paper-based microfluidic device can be used to grow plants and to hold colorimetric chemical sensors. In addition, the inherent physicochemical properties of the structured paper can be employed to simultaneously collect exudates from the roots and to bring them to the sensor at controlled times and at specific positions of the root. Using this device and the proposed extraction methodology, we can successfully extract and quantify glucose concentrations from exudates from the roots. After calibration of the sensor we could determine glucose concentrations of up to 1 mM (Figure 3C) with a LOD of 0.012 mM, which represents a slightly higher sensitivity than in our previous work (0.044 mM) (Gunatilake et al., 2021). This can be explained by the use of image analysis to quantify the signal, since the previous method used RGB data rather than HSB that is known to work better for analysis at lower concentrations. In addition, the chemical characteristics of the colorimetric sensor and the developed methodology for the extraction of the exudate over time allowed a day-by-day extraction and detection of glucose concentrations without saturating the sensor for up to 7 days.

A number of limitations of the technique arose due to the simplicity of the device. It is possible to envisage, for example, improvements of the sensor to increase the range of the detection (Mirzaei et al., 2023) or even the integration of different type of sensors such as electrochemical sensors (Cao et al., 2019; Saha et al., 2023). The time needed for extraction of the exudate was long because the sensor was not in contact with the root itself and the detection limit of the sensor. This limitation could be easily addressed, for example supplying water to the inlet using pumps and develop continuous recording of the sensor. We chose not to develop such systems because they would complexify the instrumentation needed to run experiments and put barriers to the adoption of the technologies. In addition, many changes in crop growth are studied and modelled using daily time step, *e.g.* crop growth models (Brown et al., 2018).

### Application to high throughput phenotyping studies

We used our paper-based microfluidicdevice to study the root exudation patterns of two winter wheat varieties. Although we did not observe phenotypic different and did not measure difference in growth rate, we could detect statistically significant differences in the glucose concentration of their respective root exudates (Figure 4 and 5). This result demonstrates that our techniques could easily be integrated into current high throughput root phenotyping platforms utilising paper as substrate (Le Marié et al., 2014). Since paper is a hygroscopic material, we could also envisage drastically expanding the scopes of the analyses performed by sampling the solution present at the outlet and analyse its composition using mass spectrometry techniques. Such approaches would drastically increase the ability to distinguish differences between genotypes since the detection limits of HPLC or GC-MS are in the order pM or fg, which is far higher than the sensitivity of colorimetric sensors. Complementing mass spectrometry measurements (Haase et al., 2007; Zhu et al., 2016) to continuous monitoring using colorimetric sensors would therefore significantly expand current capabilities in root phenotyping.

## Data availability

All of the generated data are available in the manuscript or in the Supplementary Information.

## Acknowledgements

This work was supported by the European Commission’s EXCELLENT SCIENCE - Marie Skłodowska-Curie Actions program, RhizoSheet MSCAIF, grant agreement number: 101028242, the MaMi project, funded by the European Union’s Horizon 2020 research and innovation programme under grant agreement No. 766007, the support from “Ministerio de Ciencia y Educación de España” under grant PID2020-120313GB-I00 / AIE / 10.13039/501100011033 and PID2020-112950RR-I00, and the Basque Government (Grant IT1633-22).

## Author contributions

Conceptualization: DP, LXD, FBL

Methodology: DP, LXD, UBG, FBL

Investigation: DP

Funding acquisition: DP, FBL, LBD, LXD

Project administration: DP, FBL

Supervision: FBL, LXD, LBD

Writing – original draft: DP, FBL

Correcting original draft: DP, FBL, LXD, LBD

## Supplementary Information (brief legends)

